# Subfield-specific Effects of Chronic Mild Unpredictable Stress on Hippocampal Astrocytes

**DOI:** 10.1101/2020.02.07.938472

**Authors:** Garima Virmani, Priyal D’almeida, Arnab Nandi, Swananda Marathe

## Abstract

Major Depressive Disorder (MDD) is a debilitating neuropsychiatric illness affecting over 20% of the population worldwide. Despite its prevalence, our understanding of its pathophysiology is severely limited, thus hampering the development of novel therapeutic strategies. Recent advances have clearly established astrocytes as major players in the pathophysiology, and plausibly pathogenesis, of major depression. In particular, astrocyte density in the hippocampus is severely diminished in MDD patients and correlates strongly with the disease outcome. Moreover, astrocyte densities from different subfields of the hippocampus show varying trends in terms of their correlation to the disease outcome. Given the central role that hippocampus plays in the pathophysiology of depression and in the action of antidepressant drugs, changes in hippocampal astrocyte density and physiology may have a significant effect on behavioral symptoms of MDD. In this study, we used Chronic Mild Unpredictable Stress (CMUS) in mice, which induces a depressive-like state, and examined its effects on astrocytes from different subfields of the hippocampus. We used S100β immunostaining to estimate the number of astrocytes per mm^2^ from various hippocampal subfields. Furthermore, using confocal images of fluorescently labeled GFAP-immunopositive hippocampal astrocytes, we quantified various morphology-related parameters and performed Sholl analysis. We found that CMUS exerts differential effects on astrocyte cell density, ramification, cell radius, surface area, and process width of hippocampal astrocytes from different hippocampal subfields. Taken together, our study reveals that chronic stress doesn’t uniformly affect all hippocampal astrocytes; but exerts its effects differentially on different astrocytic subpopulations within the hippocampus.

## Introduction

Major depression is associated with structural remodeling in various brain regions at a sub-cellular, cellular and network level (Dai et al., 2019; Nestler et al., 2002). On the other hand, antidepressant therapies rely on reversing some of these alterations by inducing structural plasticity (Castrén and Hen, 2013; Czéh et al., 2005; Santarelli et al., 2003). One of the most striking forms of structural alteration in the patients of MDD is volumetric loss as well as hypertrophy in various regions of the limbic system. In particular, major depression induces volumetric loss in the hippocampus (Bremner et al., 2002; Brown et al., 2014; Drevets, 2000; Kempton et al., 2011; Lorenzetti et al., 2009; Nestler et al., 2002; Sheline, 2003; Sheline et al., 1996). Interestingly, the volume reduction in the hippocampus shows a strong correlation with the disease outcome. In particular, a meta-analysis from 32 different studies showed that the hippocampal volume reduction was only evident among patients who experienced multiple MDD episodes or whose illness had lasted for at least 2 years (McKinnon et al., 2009). On the other hand, patients who did not fit these criteria did not show any changes in the hippocampal volume (McKinnon et al., 2009). Furthermore, in chronic or recurrent MDD, total volume of the hippocampus showed inverse correlation with the duration of the illness (Cobb et al., 2013).

Although neuronal and spine atrophy has been thought to underlie the volume reduction (Duman and Duman, 2015; Qiao et al., 2016; Sheline, 2003; Sousa et al., 2000), increasing evidence supports the involvement of astrocytes in the volumetric loss (Cobb et al., 2016; Czéh et al., 2005; Naskar and Chattarji, 2019; Rajkowska and Stockmeier, 2013; Rajkowska et al., 1999) and the pathophysiology of depression (Elsayed and Magistretti, 2015; Marathe et al., 2018). Astrocytes are profoundly affected as a result of depression, resulting in significant atrophy (Cobb et al., 2016; Cotter et al., 2001; Miguel-Hidalgo et al., 2000, 2010; Ongür et al., 1998; Rajkowska and Stockmeier, 2013; Rajkowska et al., 1999). Hippocampus is a relatively complex brain structure with 3 principal subfields viz. Dentate Gyrus (DG), CA3 and CA1 (Figure 1). In the hippocampus, the excitatory neurons are neatly arranged in thin layers of cells, such as the granule cell layer of the dentate gyrus and the principal cell layers of CA3 and CA1 (Figure 1). These neuronal layers are relatively devoid of astrocytic cell bodies. However, the regions of the hippocampus that are rich in synapses, such as the molecular layer of the DG, harbor dense populations of astrocytes. Given their location close to the synapses, any detrimental change in their morphology may have a large impact on hippocampus-dependent behaviors. It appears that the atrophy in astrocytes is not uniform and varies between different hippocampal subfields (Cobb et al., 2013, 2016; Willard et al., 2013). In particular, region-specific changes in astrocytic density that may correlate with behavioral symptoms and the history of antidepressant use are of immense interest. Intriguingly, it was shown that in the absence of antidepressant treatment, astrocytic density in the hippocampal hilar region was reduced (Cobb et al., 2016). On the other hand, the depressed patients taking antidepressant medication did not show any such decrease in hilar astrocyte density (Cobb et al., 2016). The same study also showed an inverse correlation between area fraction of GFAP immunoreactivity in the CA2/3 region, and the duration of depression in suicide victims (Cobb et al., 2016). Additionally, the volume fraction of GFAP immunoreactivity was selectively decreased in the DG in females with depression, but not in males (Cobb et al., 2016). A study in depressed monkeys showed that depression was associated with reduced astrocyte density in the DG and CA1 of the anterior hippocampus (Willard et al., 2013). These results suggest that prolonged depression may exert differential effects on the astrocytes from different subfields of the hippocampal formation.

**Figure 1:**
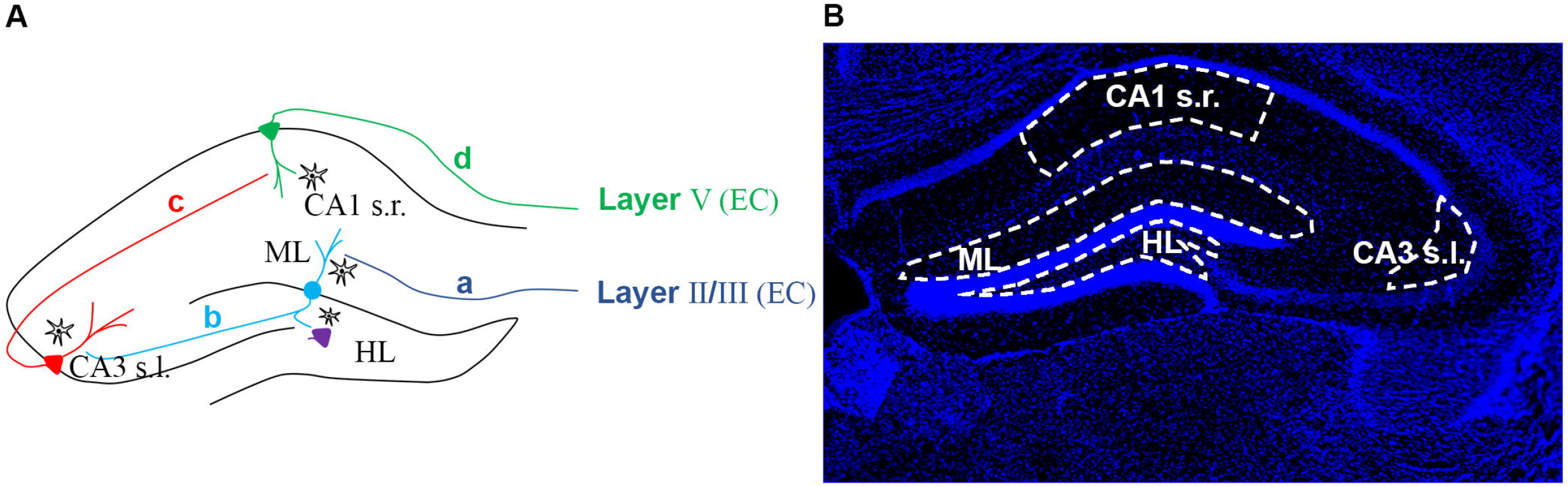
Hippocampal circuitry and ROI selection. **(A)** Hippocampus is characterized by trisynaptic loop comprising of: (a) inputs through perforant pathway originating from layers II/III of the entorhinal cortex (EC), forming synapses with dentate gyrus (DG) granule cells in the molecular layer (ML). (b) The mossy fiber pathway originating from the DG granule cells and forming synapses with mossy cells in the hilus (HL) as well as pyramidal neurons of the CA3 in the CA3 *stratum lucidum* (CA3 s.l.). (c) The schaffer collaterals originating from CA3 pyramidal neurons and forming synapses with the CA1 pyramidal neurons in the *stratum radiatum* of the CA1 (CA1 s.r.). (d) Output from the CA1 pyramidal neurons goes back to the layer V of the EC. **(B)** Image of the entire hippocampus captured through a 4X objective showing DAPI-positive nuclei. The regions of interest (ROIs) used throughout this study have been outlined with white dotted lines. The ROIs were located in regions rich in synapses and devoid of the principal excitatory neurons of the hippocampus.

How depression may affect astrocytes from different hippocampal subfields differently is not clear. Mechanistic understanding of this phenomenon of region-specific differential effects would require this to be studied in simpler and more tractable model system. Astrocytic atrophy has been studied in animal models of depression (Czéh et al., 2005; Jang et al., 2008; Musholt et al., 2009; Naskar and Chattarji, 2019). However, these studies haven’t compared the astrocytes across various hippocampal subfields along the rostro-caudal extent of the hippocampus. In this study, we sought to address whether such subfield-specific changes in astrocyte density and morphology can be found in the hippocampus of mouse model of depression as well.

In this study, we subjected C57BL/6J mice to chronic mild unpredictable stress (CMUS), that induces depressive-like state and studied the changes in the astrocyte density and morphology across different hippocampal subfields. We found subfield-specific effects on cell density and on morphology-related parameters.

We believe that these results would prove useful to understand the mechanisms underlying diversity and differential vulnerability of hippocampal astrocytes in the pathophysiology of depression.

## Materials and Methods

### Animals

All experiments were performed on C57BL/6J mice bred in IISc Central Animal Facility. Mice were group-housed and maintained on 12/12-hour light/dark cycle with access to food and water *ad libitum*. Adult male mice (25-30 g, 4-5 months old) were used for all experiments. All procedures were carried out in accordance with the protocols approved by Institutional Animal Ethics Committee (IAEC), Indian Institute of Science. All efforts were made to reduce the number of animals by using the principle of 3R’s.

### CMUS

Mice undergoing CMUS were transported to another room 30 minutes prior to the onset of stress. Control animals remained in the holding room to avoid exposure to stress-related auditory and olfactory stimuli from the CMUS-treated mice. 30 minutes after the completion of each stressor, the mice were brought back to the holding room. 1-2 stressors were administered every day at random time intervals with a gap of at least 3 hours between successive stressors (Supplementary Table 1). Following stressors were used: Overcrowding (3-4 mice kept in 1 litre glass beaker for 3 hours), Cage Tilt (Home cages were tilted at 45 degrees for 12 hours overnight), Cold Exposure (Cages were kept on a layer of ice for 45 minutes), Wet Bedding (2 cm deep water was added to home cages for 3 hours), Restraint (mice were restrained in a 50ml conical plastic tube with a hole in the bottom to facilitate breathing), Food and Water Deprivation (Food and water bottles were removed for 12 hours overnight), Shaker Stress (Cages were kept on a shaker (Tarsons Dancing Shaker MC-02 Cat no. 3080) run at 50 rpm for 3 hours), Tone (10 kHz tone was played at about 80 Db for 12 hours overnight), White Noise (White noise was played in the band of 0-20 kHz at about 80 Db for a period of 12 hours overnight). Specific schedule used in this study is summarized in the supplementary table 1 (Supplementary Table 1).

### Forced Swim Test (FST)

FST was a part of the CMUS paradigm and served the dual purpose of being one of the stressors and also as a test of behavioral analysis. Control mice underwent FST on day 0 and Day 19. In brief, mice were allowed to swim in a 2-liter glass beaker for 5 minutes and were video recorded from the side. Immobility was scored manually. Mice were deemed to be immobile when there was a lack of any discernible movement of the limbs, other than what is required to stay afloat. Time points at which mice switched from mobility to immobility and *vice versa* were recorded and used for further analysis. Immobility on day 0 was compared with that on day 19 using three parameters: total time immobile (seconds), latency to first continuous bout of immobility of ≥ 15 seconds (seconds) and the duration of the longest bout of immobility (seconds). Tail suspension test was a part of the CMUS paradigm, but it was not used for monitoring behavioral changes since mice showed extensive tail-climbing behavior rendering the analysis inaccurate.

### Tissue preparation

24 hours following the last stressor, mice were anaesthetized using isoflurane and were sacrificed by transcardial perfusion with Phosphate buffered saline (PBS) followed by 4% paraformaldehyde (PFA). Serial floating coronal sections (40μm thick) were cut using a cryostat (Leica, CM 1850) and were stored at −20°C in a freezing mixture (2:1:1 mixture of 0.1M Phosphate Buffer (PB): Ethylene Glycol: Glycerol) until further use.

### Immunohistochemistry and imaging

Immunohistochemistry protocol was based on those described earlier (Marathe et al., 2015; Yanpallewar et al., 2010). In brief, the sections were removed from the freezing mixture and washed 3 times with PB. Sections were then blocked for 1 hour with 10% normal donkey serum + 3% Bovine Serum Albumin (BSA) + 0.3% TritonX100 in 0.1M PB. Sections were incubated overnight at 4°C with primary antibodies (chicken pAb anti-GFAP; 1:1000; Novus, NBP1-05198 and rabbit pAb anti S100b; 1:1000, Synaptic systems287003). On the second day, the sections were washed 3 times with PB and incubated for 2 hours at room temperature with secondary antibodies (goat anti chicken Alexa Flour 594, abcam, ab150172; and donkey anti rabbit Alexa Flour 488, abcam, ab150073). After incubation, sections were washed 3 times in PB and were mounted on a glass slide in the mounting medium with DAPI (Abcam ab 104139). The slides were imaged at 20X on a Zeiss LSM 880 airyscan confocal microscope.

### Cell Density Analysis

For calculating the density of astrocytes, 3 sections per mouse were used per data point. 3-4 mice were used per group. z-stacks from the confocal images of S100β-positive cells were flattened with maximum intensity projection. A region of interest (ROI) was selected encompassing the given hippocampal subfield. We restricted our ROIs in CA3 largely to *stratum lucidum* and in CA1 to *stratum radiatum*. We calculated the number of S100β-positive nuclei and divided it by the area of the selected ROI, using ImageJ. The numbers were then converted to the number of astrocytes/mm^2^ by multiplying by an appropriate factor. In order to estimate the astrocyte density across the rostro-caudal axis, the data was generated from sections from different bregma levels. The sections between bregma levels - 1.46 to −1.94 were labelled as rostral, −2.06 to −2.46 as intermediate and sections beyond −2.54 were labelled as caudal.

### Morphological Analysis

Morphological analysis was done using Ghosh Lab Sholl analysis plugin in ImageJ. In brief, maximum intensity projection of confocal images containing GFAP-stained astrocytes was generated using ZEN black edition 3.0 SR and saved in TIFF format. The 2-D images were then binarized in ImageJ. We then used “analyze particles” tool to eliminate the background using size exclusion. Single cells were cropped out and surface area was measured by pixel count. Next, we filled holes in the thresholded image using “fill holes” tool in ImageJ. Following this step, the cells were skeletonized using skeletonize plugin in ImageJ to generate cells with single pixel-wide processes. The pixel count of the skeletonized cell was calculated to estimate the total process length. Surface area of every cell was divided by its total process length to generate the average process width for that cell. The centroid for Sholl analysis was selected based on the morphology of the original GFAP-stained images. Sholl analysis plugin from Ghosh lab was then used to generate Sholl analysis vectors. A 5-period moving average was calculated for every cell to generate smooth curves for Sholl analysis, as polynomial regressions were deemed unsuitable for our datasets. Approximately 60-150 cells (from 4 sections per mouse) from 3-4 mice per group were used for Sholl analysis per region per group. The results were represented as mean ± S.E.M. of all cells. Area under the Sholl curve for every cell was calculated by taking the sum of the Sholl vector. Enclosing radius of each cell was calculated by the length of the respective Sholl vector after removing the zeros.

### Statistical Analysis

All statistical analyses were performed using GraphPad Prism 8.0.2. Results are expressed as mean ± SEM for Sholl analysis and FST analysis; and as box and whiskers plots for 2 group comparisons. Boxes represent the 25^th^ to 75^th^ percentile whereas whiskers represent the minimum and maximum value in the corresponding dataset, with median represented as a horizontal line across the box and mean as “+”. Requisite sample sizes were determined using power analysis. Cell count comparisons between two groups were made using unpaired student’s t-test. The analysis of astrocyte densities along the rostro-caudal axis was done using mixed effects model to account for the interactions between the bregma levels and stress exposure. This was followed by corrected Sidak’s *post-hoc* test. For 2 group comparisons of morphology-related parameters, normality of the datasets was determined using Kolmogorov-Smirnov normality test with Dallal-Wilkinson-Lilliefor approximation. The distribution was deemed to be normal (Gaussian) at the alpha of 0.05. Two group comparisons were made using nested t-test to determine statistical significance. Nested t-test was used to account for the nested design in experimental samples, where analyzed cells came from different biological replicates (mouse identity). For Sholl analysis, the data was analyzed using two-way repeated measures ANOVA with Geisser-Greenhouse correction to account for possible unequal variances (Wilson et al., 2017). This was followed by corrected Sidak’s *post-hoc* test.

## Results

### Effects of CMUS on astrocyte density across hippocampal subfields

To study the effects of stress on astrocyte density and morphology, we subjected mice to 21 days of CMUS as described in materials and methods section (Supplementary Table 1). The emergence of depressive-like symptoms was confirmed using FST (Supplementary Figure 1). The CMUS-treated mice showed trend towards an increase in time spent immobile from day 0 to day 19 (Day 0: 51.82 ± 5.77, Day 19: 74.15 ± 4.26) (p=0.078, paired student’s t-test) (Supplementary Figure 1 D), while the control mice did not show any change (Day 0: 61.95 ± 1.53, Day 19: 63.83 ± 3.16) (p=0.69, paired student’s t-test) (Supplementary Figure 1 A). The CMUS-treated mice also showed an increase in the duration of the longest bout of immobility (Day 0: 22.33 ± 3.84, Day 19: 39.67 ± 7.06) (p=0.03, paired student’s t-test) (Supplementary Figure 1 E) and a decrease in the latency to a continuous bout of immobility of 15 seconds or longer (Day 0: 122.70 ± 34.72, Day 19: 18.33 ± 14.38) (p=0.04, paired student’s t-test) (Supplementary Figure 1 F). On the other hand, the control mice did not show any change in these two parameters (Duration of the longest bout: Day 0: 48.75 ± 10.48, Day 19: 27.50 ± 3.66; p=0.12, paired student’s t-test) (Latency to first immobility bout of ≥ 15 seconds: Day 0: 122.30 ± 12.76, Day 19: 77.75 ± 45.36; p=0.44, paired student’s t-test) (Supplementary Figure 1 B, C).

In all the hippocampal subfields that we analyzed from C57BL/6J mice, we found a near complete overlap between cells expressing S100β and GFAP (Supplementary Figure 2). Hence, we used the astrocytic markers S100β and GFAP to analyze astrocyte density and morphology respectively in our study.

In order to assess the effects of CMUS on the astrocyte density, we estimated the number of astrocytes per mm^2^ as described in the materials and methods section. We did not find a difference in astrocyte density in the molecular layer of the DG (Control: 101.00 ± 5.87, CMUS: 88.18 ± 4.13) (p=0.096, unpaired student’s t-test) (Figure 2A). Astrocyte density analysis in the hilar region of the DG revealed no significant difference between the groups in the hilus (Control: 222.40 ± 21.34, CMUS: 177.40 ± 12.89) (p=0.11, unpaired student’s t-test). Overall, the astrocyte density in the hilus was approximately twice that in the molecular layer, which was consistent with previous studies (Figure 2A and 2B). In *stratum lucidum* region of CA3, we did not find any difference between CMUS-treated mice and control mice (Control: 126.20 ± 9.95, CMUS: 124.80 ± 8.84) (p=0.932, unpaired student’s t-test) (Figure 2C). We then quantified the astrocyte density in the *stratum radiatum* region of CA1. Again, we did not find a statistically significant difference in the number of astrocytes per mm^2^ in CA1 (Control: 103.40 ± 7.96, CMUS: 85.52 ± 5.88) (p=0.08, unpaired student’s t-test) (Figure 2D). Taken together, these results indicate that CMUS does not induce a statistically significant change in the number of astrocytes per mm^2^ in any of the hippocampal subfields that we tested when sections are taken spanning the entire rostro-caudal extent.

**Figure 2:**
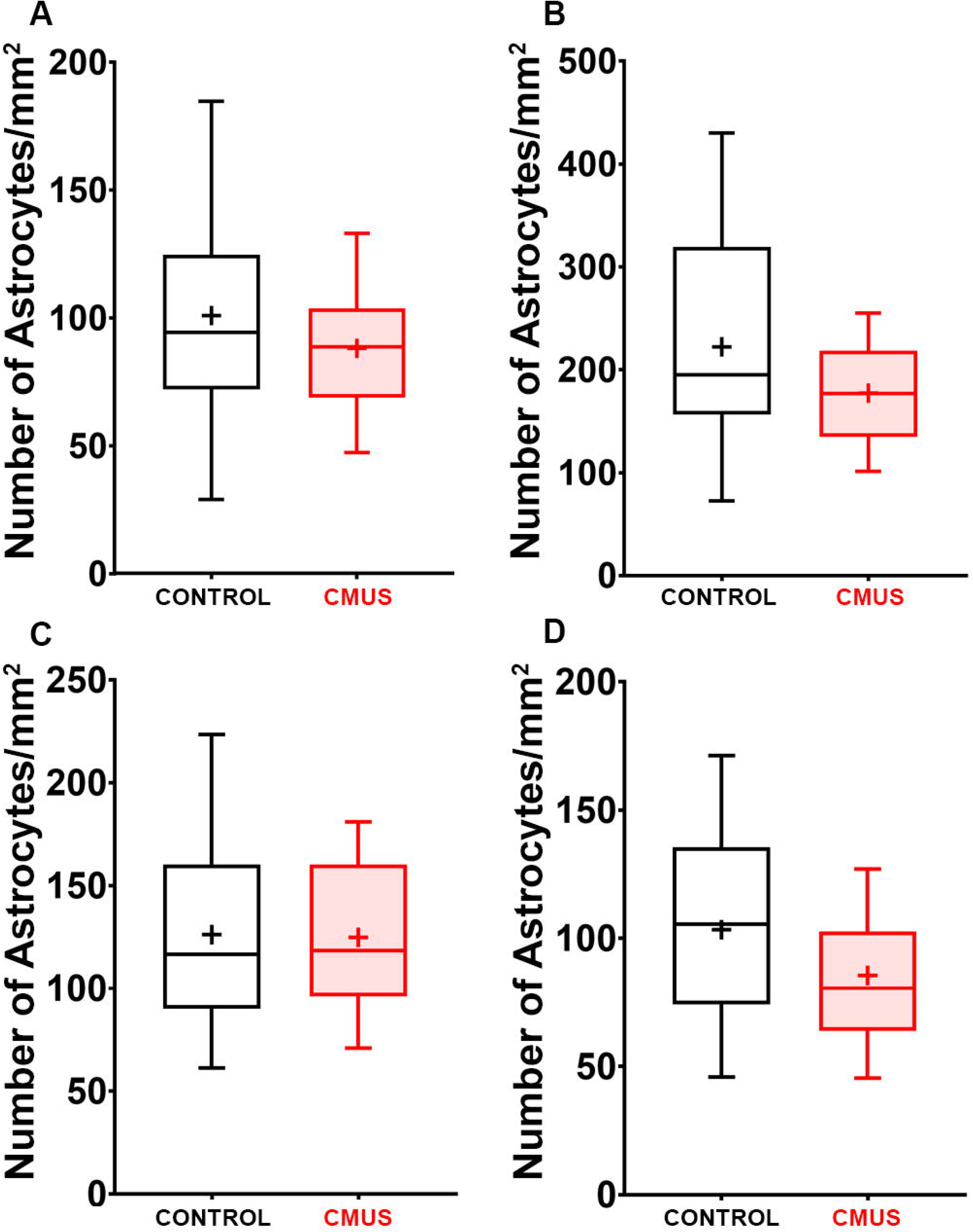
Effects of CMUS on astrocyte densities in different hippocampal subfields. Mice were subjected to 21 days of Chronic Mild Unpredictable Stress (CMUS), while the controls were handled similarly without exposure to stress. S100β immunostained astrocytes were imaged on a confocal microscope. ROI was marked on 2-D maximum intensity projections encompassing the desired subfield and the number of cells counted was divided by the area of the ROI. The data was represented and analyzed as the number of astrocytes/mm^2^. We did not find statistically significant difference in the molecular layer of the DG **(A)**, hilus **(B)**, *stratum lucidum* of the CA3 **(C)** or *stratum radiatum* of the CA1 **(D)**. n=3-4 mice per group. Data represented as box and whisker’s plot. All comparisons made using unpaired student’s t-test.

### Effects of CMUS on astrocyte density in rostral, intermediate and caudal regions of the hippocampus

Next, we sought to address whether the astrocytic density is differentially affected between the rostral, intermediate and the caudal regions of different hippocampal subfields. We took sections from rostral (Bregma level −1.46 to −1.94), intermediate (Bregma level −2.06 to −2.46) and caudal (Bregma level −2.54 to −2.92) to estimate the astrocytic density. In the molecular layer of the DG, we did not find any difference between the two groups in rostral (Control: 104.30 ± 9.24, CMUS: 85.93 ± 5.85) (p=0.29), intermediate (Control: 83.53 ± 9.51, CMUS: 85.84 ± 7.75) (p=0.99) or caudal (Control: 110.10 ± 10.43, CMUS: 93.21 ± 8.60) (p=0.53) (mixed effects analysis to assess interactions between rostro-caudal level and stress treatment, followed by corrected Sidak’s *post-hoc* test) (Figure 3E). In the rostral region, we saw a significant decrease in the astrocyte density (Control: 340.50 ± 17.75, CMUS: 198.90 ± 18.67) (p=0.0006), whereas there was no significant difference in the astrocyte density in the intermediate (Control: 124.3 ± 18.80, CMUS: 137.80 ± 15.58) (p=0.93) and the caudal (Control: 192.60 ± 9.29, CMUS: 194.50 ± 25.51) (p=0.99) hilus (mixed effects analysis to assess interactions between rostro-caudal level and stress treatment, followed by corrected Sidak’s *post-hoc* test) (Figure 3A, B, F). Furthermore, we assessed if CMUS affects the astrocyte morphology in the *stratum lucidum* of the CA3 along the rostro-caudal axis. However, we did not find any statistically significant difference in the astrocyte density in rostral (Control: 170.80 ± 15.20, CMUS: 144.20 ± 14.54) (p=0.54), intermediate (Control: 89.70 ± 8.03, CMUS: 109.80 ± 8.24) (p=0.30) and caudal (Control: 119.00 ± 13.12, CMUS: 116.50 ± 19.15) (p=0.99) regions of CA3 (mixed effects analysis to assess interactions between rostro-caudal level and stress treatment, followed by corrected Sidak’s *post-hoc* test) (Figure 3G). In the *stratum radiatum* region of the CA1, we found a significant decrease in the rostral part of CA1 (Control: 129.10 ± 12.45, CMUS: 82.64 ± 9.52) (p=0.03), whereas no differences were observed in the intermediate (Control: 75.47 ± 11.84, CMUS: 82.53 ± 11.44) (p=0.96) and caudal (Control: 98.29 ± 10.35, CMUS: 92.57 ± 11.00) (p=0.97) regions (mixed effects analysis to assess interactions between rostro-caudal level and stress treatment, followed by corrected Sidak’s *post-hoc* test) (Figure 3C, D, H). Together, these results indicate that the astrocyte density is decreased only in rostral hilus and rostral CA1 as a result of CMUS, indicating region-specific alterations.

**Figure 3:**
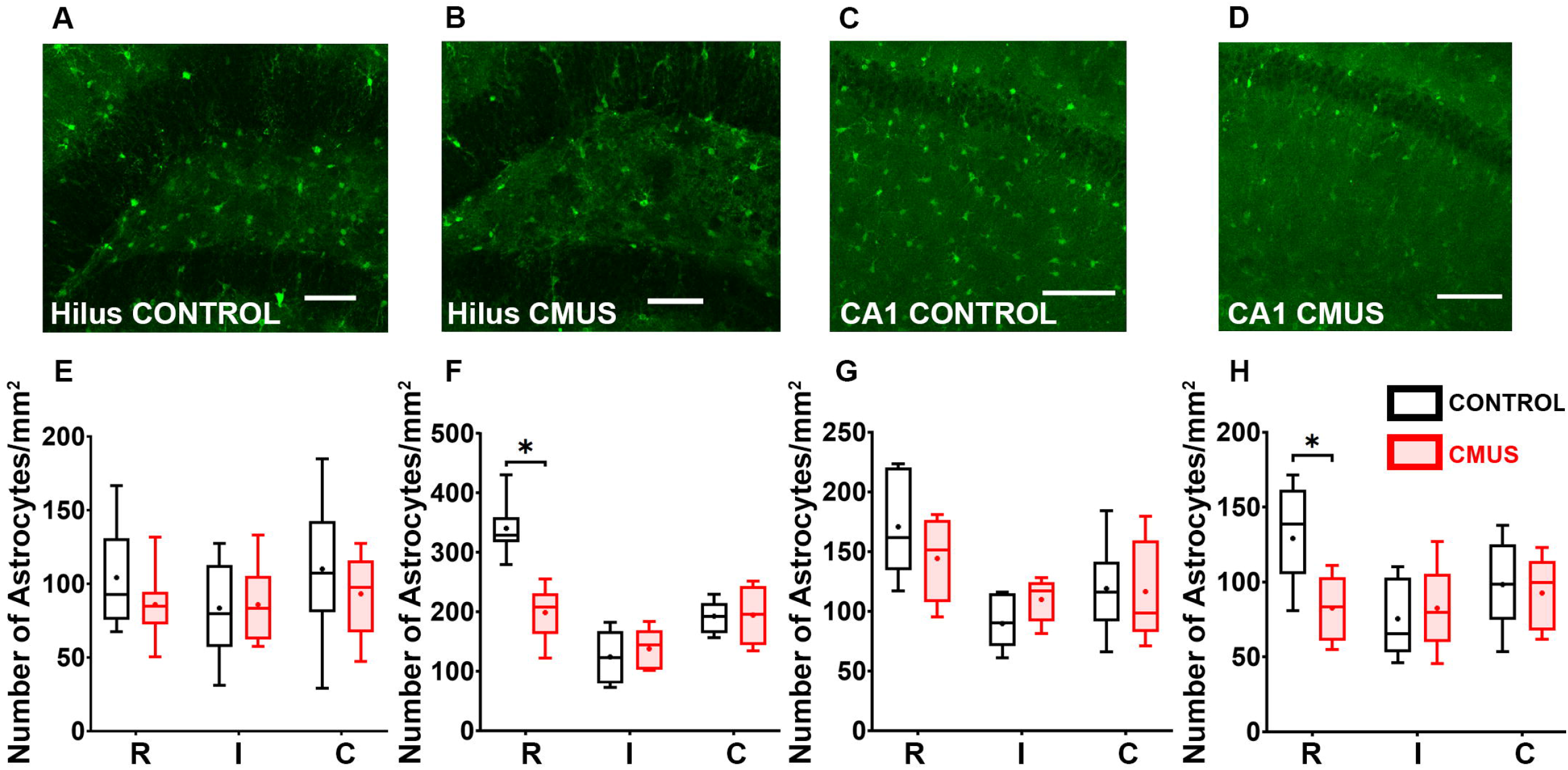
Effects of CMUS on astrocyte densities across the rostro-caudal axis in different hippocampal subfields. Mice were subjected to 21 days of Chronic Mild Unpredictable Stress (CMUS), while the controls were handled similarly without exposure to stress. S100β immunostained astrocytes were imaged on a confocal microscope. ROI was marked on 2-D maximum intensity projections encompassing the desired subfield and the number of cells counted was divided by the area of the ROI. The data was represented and analyzed as the number of astrocytes/mm^2^. The data was binned into 3 categories viz. Rostral (R), Intermediate (I) and Caudal (C) depending on the bregma levels (see materials and methods). We saw a statistically significant decrease in the astrocyte density in the rostral hippocampus of the hilus **(A, B, F)** and the *stratum radiatum* of the CA1 **(C, D, H)**. All other comparisons yielded no statistically significant differences **(E, F, G, H)**. n=3-4 mice per group. Data represented as box and whisker’s plot. * represents p<0.05. All comparisons made with mixed-effects model to study the differential effects of dorso-ventral position and stress treatment, followed by Sidak’s *post-hoc* test. Scale bars in **(A, B, C, D)** are 100μm.

### Effects of CMUS on the morphology of astrocytes in the molecular layer of the DG

Since we found that CMUS exerts its effects on astrocyte density in the rostral parts, and that intermediate and caudal parts seem to be spared in our analysis, we next addressed if CMUS affects the morphology of astrocytes in the rostral sections. The morphological analysis on GFAP-stained images of astrocytes was done as explained in the materials and methods section, and in figure 4 (Figure 4). To assess the effects of CMUS on astrocytic branching on molecular layer astrocytes, we performed Sholl analysis. We found a significant decrease in the ramification along most of the radius of cells in CMUS-treated mice as compared to the control mice (two-way repeated measures ANOVA followed by Sidak’s *post-hoc* test) (Figure 5A). We then compared the area under the Sholl analysis curve between the controls and CMUS-treated mice and found a significant decrease (Control: 85.99 ± 2.04, CMUS: 73.12 ± 1.48) (p=0.006, nested t-test), suggesting a robust decline in ramifications (Figure 5B). We then studied the enclosing radius and found a significant decrease in the CMUS-treated mice as compared to the control mice (Control: 27.07 ± 0.47, CMUS: 25.02 ± 0.47) (p=0.04, nested t-test) (Figure 5C). Next, we calculated the total surface area covered by astrocytes from the thresholded maximum intensity projected images and found a significant decrease in the CMUS-treated group as compared to the control mice (Control: 214.30 ± 5.96, CMUS: 185.40 ± 4.13) (p=0.0002, nested t-test) (Figure 5D). However, we did not find any difference in the average width of astrocytic processes between the control and CMUS-treated mice (Control: 2.35 ± 0.02, CMUS: 2.44 ± 0.03) (p=0.12, nested t-test) (Figure 5E).

**Figure 4:**
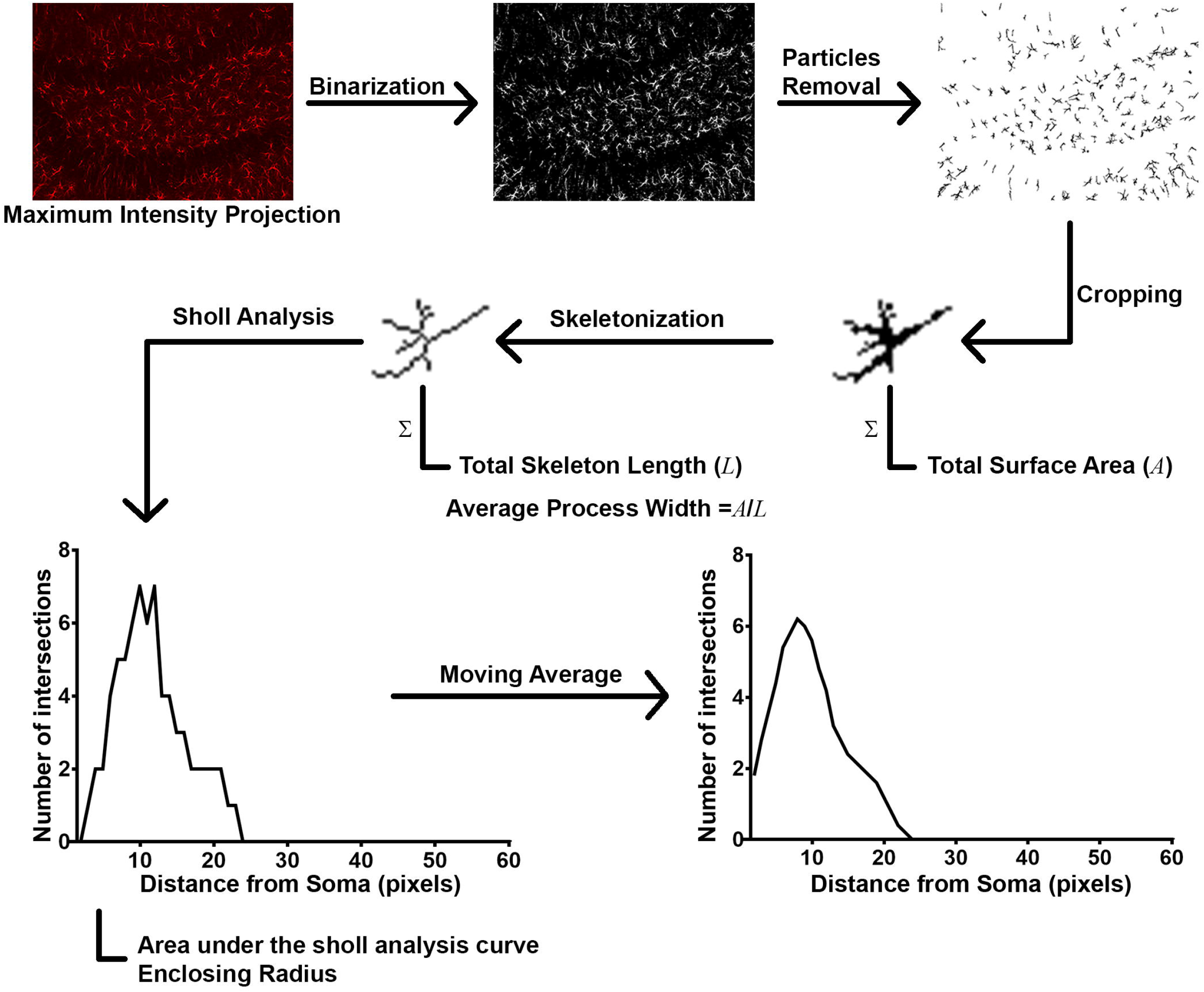
Image analysis pipeline: Images of GFAP-stained hippocampal sections were captured on a confocal microscope and maximum intensity projection was generated. The images were then binarized by using global dynamic thresholding. The background was eliminated using size exclusion. Single cells were cropped out and surface area was measured by pixel count. Next, the cells were skeletonized to generate cells with single pixel-wide processes. The pixel count of the skeletonized cell was calculated to estimate the total process length. Surface area of every cell was divided by its total process length to generate the approximate average process width for that cell. Skeletonized images of astrocytes were used to generate Sholl analysis vectors. A 5-period moving average was calculated for every cell to generate smooth curves for Sholl analysis.

**Figure 5:**
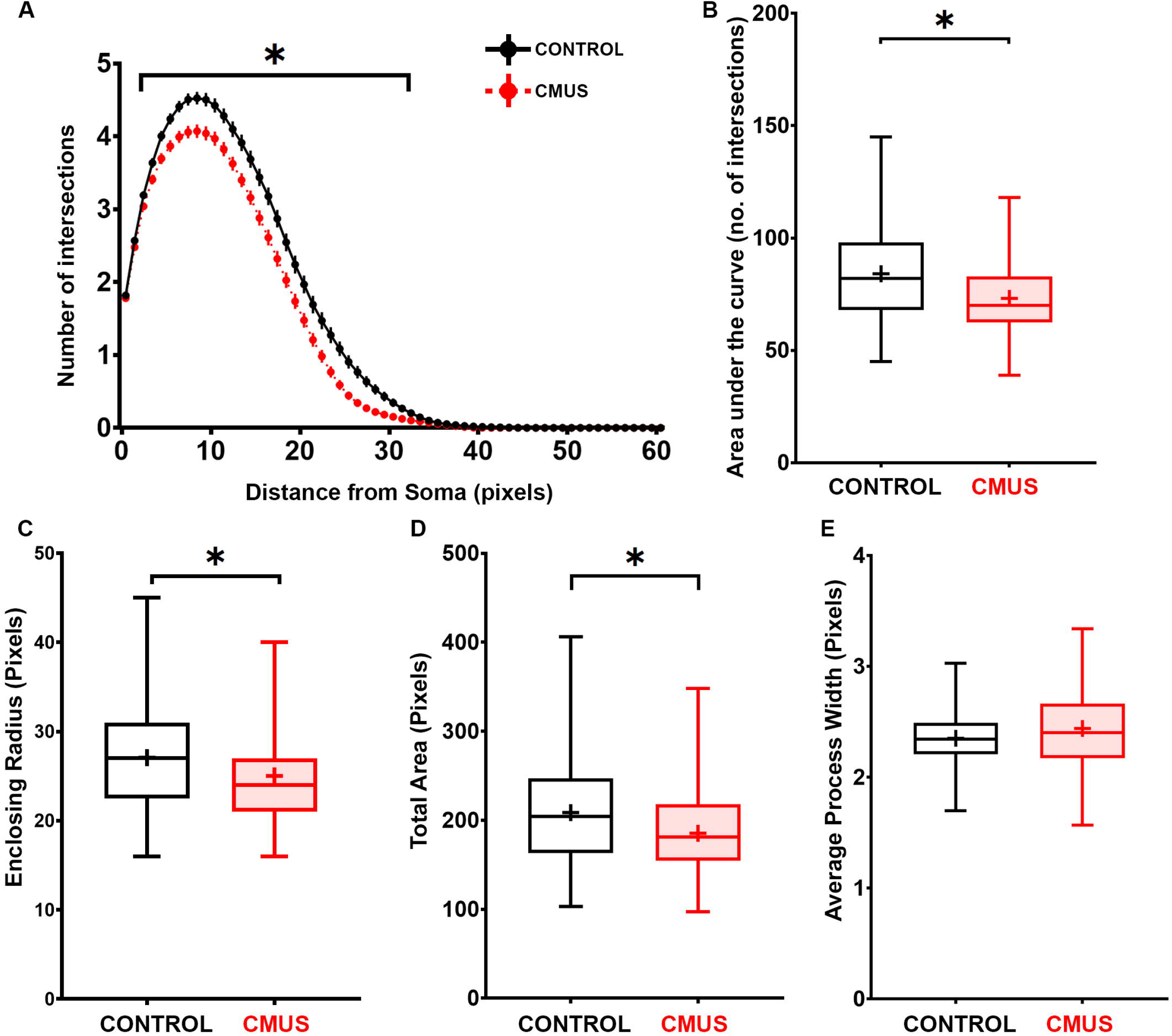
CMUS affects the morphology of astrocytes in the molecular layer of the DG. Mice were subjected to 21 days of Chronic Mild Unpredictable Stress (CMUS), while the controls were handled similarly without exposure to stress. GFAP immunostained astrocytes were imaged on a confocal microscope and the images of astrocytes from the molecular layer of the DG (ML) were used for morphological analysis as shown in Figure 4. Line plot depicting Sholl analysis reveals a significant decrease in the ramification of astrocytes in CMUS-treated mice as compared to the control mice **(A)**. Box and whiskers plots show a significant decrease in the area under the Sholl curve **(B)**, Enclosing Radius **(C)**, and Total projected surface area **(D)** for astrocytes from CMUS-treated mice as compared to the controls. However, no statistically significant difference was seen in the average process width between the two groups **(E)**. n= (control: 145 cells from 4 mice, CMUS: 121 cells from 3 mice). Data represented as mean ± SEM for Sholl analysis and as box and whisker’s plot for two group comparisons. For Sholl analysis, * represents p<0.05, Two-way repeated measures ANOVA followed by Sidak’s *post-hoc* test. For Two group comparisons, * represents p<0.05, nested t-test.

### Effects of CMUS on the morphology of astrocytes in the hilus

Next, we assessed the effects of CMUS on astrocytic branching on hilar astrocytes by Sholl analysis. We found a significant decrease in the ramification, but only along a small portion of the proximal segment of astrocytic branches in CMUS-treated mice as compared to the control mice (two-way repeated measures ANOVA followed by Sidak’s *post-hoc* test) (Figure 6A). However, we found no difference in the area under the Sholl analysis curve between the controls and CMUS-treated mice (Control: 55.49 ± 1.49, CMUS: 51.44 ± 1.71) (p=0.18, nested t-test) (Figure 6B), suggesting only a marginal change in branching at a small portion of proximal branches (Figure 5A, B). We also found no statistically significant difference between CMUS-treated mice and control mice in terms of the enclosing radius (Control: 19.30 ± 0.37, CMUS: 18.52 ± 0.50) (p=0.42, nested t-test) (Figure 6C) and total surface area (Control: 130.50 ± 3.71, CMUS: 128.50 ± 4.77) (p=0.95, nested t-test) (Figure 6D). Interestingly, we found a small (7.56 %), but statistically significant (p=0.001, nested t-test) increase in average process width in astrocytes of CMUS-treated mice as compared to the controls (Control: 2.29 ± 0.03, CMUS: 2.46 ± 0.05) (Figure 6E).

**Figure 6:**
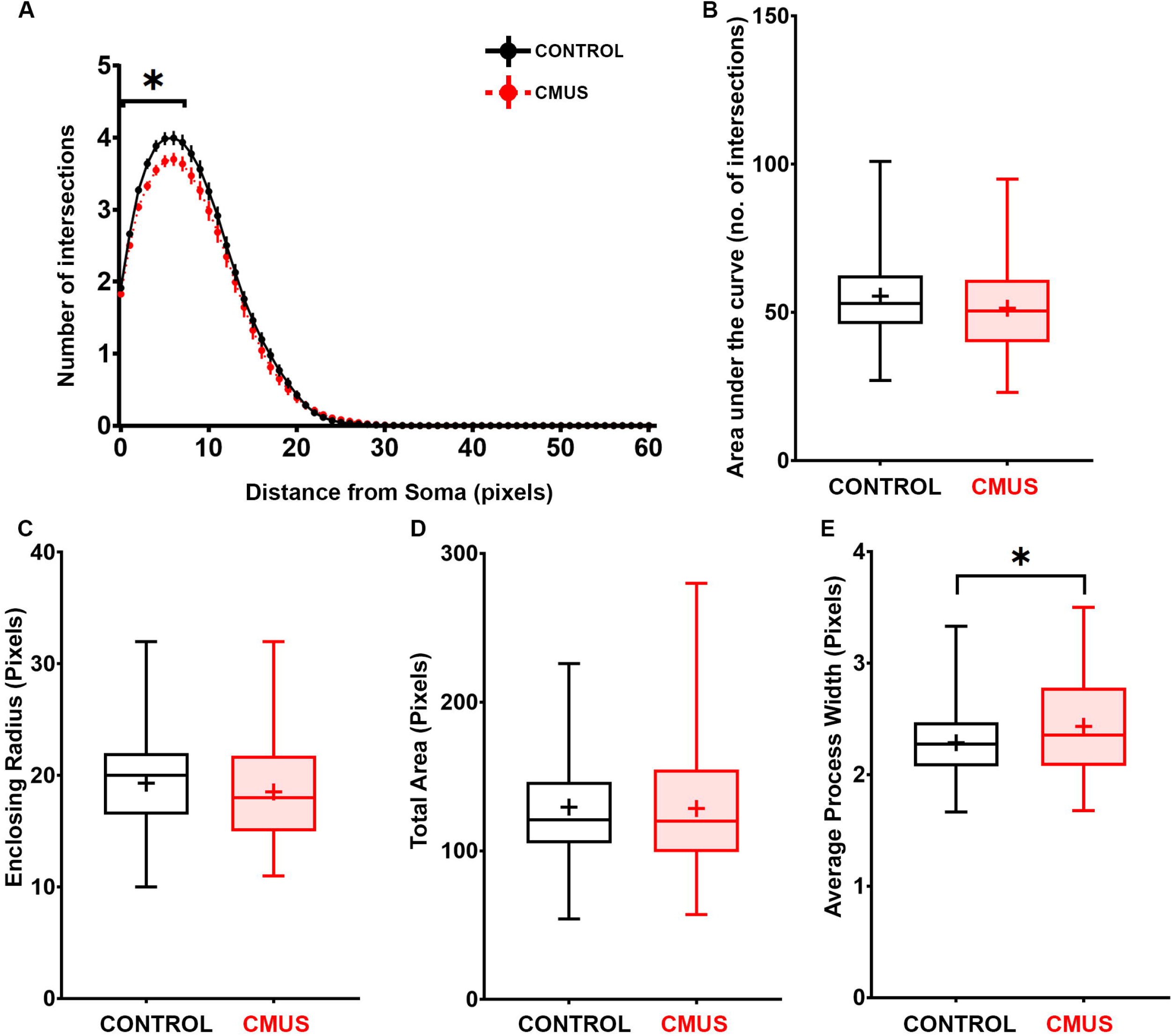
Effects of CMUS on the morphology of astrocytes in the hilus. Mice were subjected to 21 days of Chronic Mild Unpredictable Stress (CMUS), while the controls were handled similarly without exposure to stress. GFAP immunostained astrocytes were imaged on a confocal microscope and the images of astrocytes from the hilar region of the DG were analyzed for morphological analysis as shown in Figure 4. Line plot depicting Sholl analysis reveals a significant decrease in the ramification only in the proximal segment of astrocytes in CMUS-treated mice as compared to the control mice **(A)**. Box and whiskers plots show no significant change in the area under the Sholl curve **(B)**, Enclosing Radius **(C)**, and Total projected surface area **(D)** for astrocytes from CMUS-treated mice as compared to the controls. However, a small (7.56%) but statistically significant increase was seen in the average process width in the hilar astrocytes of CMUS-treated mice as compared to the control mice **(E)**. n= (control: 109 cells from 4 mice, CMUS: 84 cells from 3 mice). Data represented as mean ± SEM for Sholl analysis and as box and whisker’s plot for two group comparisons. For Sholl analysis, * represents p<0.05, Two-way repeated measures ANOVA followed by Sidak’s *post-hoc* test. For Two group comparisons, * represents p<0.05, nested t-test.

### Effects of CMUS on the morphology of astrocytes in the stratum lucidum of CA3

In order to study the effects of CMUS on branching on astrocytes from the *stratum lucidum* of the CA3, Sholl analysis was performed. We found a statistically significant decrease in the ramification, but only in a small portion of distal branches of astrocytes in CMUS-treated mice as compared to the control mice (Figure 7A). At the other segments, we did not find any change in ramification (Figure 7A) (two-way repeated measures ANOVA followed by Sidak’s *post-hoc* test). We also examined other morphology-related parameters in these astrocytes from the *stratum lucidum* of CA3. However, we did not find a statistically significant difference in the area under the Sholl curve (Control: 85.14 ± 3.36, CMUS: 77.92 ± 2.82) (p=0.21, nested t-test) (Figure 7B), enclosing radius (Control: 28.03 ± 0.96, CMUS: 27.16 ± 0.89) (p=0.62, nested t-test) (Figure 7C) or in total area of the cells (Control: 214.30 ± 9.95, CMUS: 200.50 ± 9.26) (p=0.46, nested t-test) (Figure 7D) between the control mice and the CMUS-treated mice. Additionally, we did not find a statistically significant difference in the average process width between the CA3 astrocytes from control mice and the CMUS-treated mice (Control: 2.41 ± 0.04, CMUS: 2.51 ± 0.06) (p=0.17, nested t-test) (Figure 7E).

**Figure 7:**
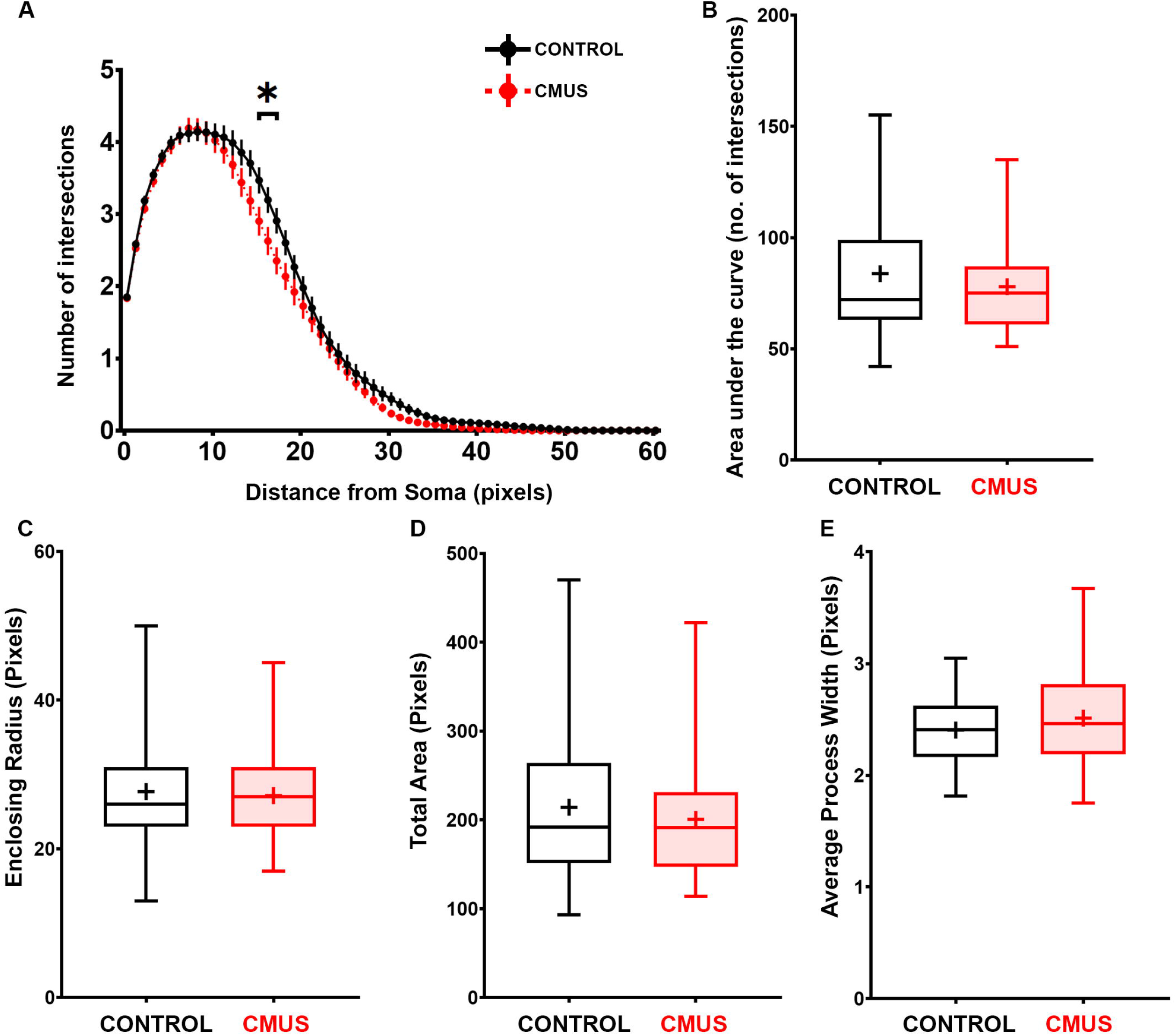
Effects of CMUS on the morphology of astrocytes in the stratum lucidum of CA3. Mice were subjected to 21 days of Chronic Mild Unpredictable Stress (CMUS), while the controls were handled similarly without exposure to stress. GFAP immunostained astrocytes were imaged on a confocal microscope and the images of astrocytes from the *stratum lucidum* region of the CA3 were analyzed for morphological analysis as shown in Figure 4. Line plot depicting Sholl analysis reveals a significant decrease in the ramification only in a small portion of the distal segment of astrocytes in CMUS-treated mice as compared to the control mice **(A)**. Box and whiskers plots show no significant change in the area under the Sholl curve **(B)**, Enclosing Radius **(C)**, Total projected surface area **(D)** and Average process width **(E)** for astrocytes from CMUS-treated mice as compared to the controls. n= (control: 72 cells from 4 mice, CMUS: 51 cells from 3 mice). Data represented as mean ± SEM for Sholl analysis and as box and whisker’s plot for two group comparisons. For Sholl analysis, * represents p<0.05, two-way repeated measures ANOVA followed by Sidak’s *post-hoc* test. Two group comparisons were analyzed with nested t-test.

### Effects of CMUS on the morphology of astrocytes in the stratum radiatum of CA1

We further investigated the effects of CMUS on astrocytic branching in the *stratum radiatum* of the CA1 using Sholl analysis. However, we did not find any difference in ramification between astrocytes from CMUS-treated mice as compared to the control mice (two-way repeated measures ANOVA followed by Sidak’s *post-hoc* test) (Figure 8A). Furthermore, we did not find a statistically significant difference in the area under the Sholl curve (Control: 81.74 ± 2.59, CMUS: 73.74 ± 2.62) (p=0.25, nested t-test) (Figure 8B), enclosing radius (Control: 28.19 ± 0.81, CMUS: 26.27 ± 0.89) (p=0.18, nested t-test) (Figure 8C) or in total area of the cells (Control: 197.40 ± 7.47, CMUS: 177.50 ± 8.30) (p=0.30, nested t-test) (Figure 8D). We also did not find any difference in the average process width between the CA1 *stratum radiatum* astrocytes from control mice and the CMUS-treated mice (Control: 2.33 ± 0.03, CMUS: 2.31 ± 0.05) (p=0.76, nested t-test) (Figure 8E). These results indicate that the astrocytes from CA1 *stratum radiatum* are not affected morphologically in our CMUS paradigm.

**Figure 8:**
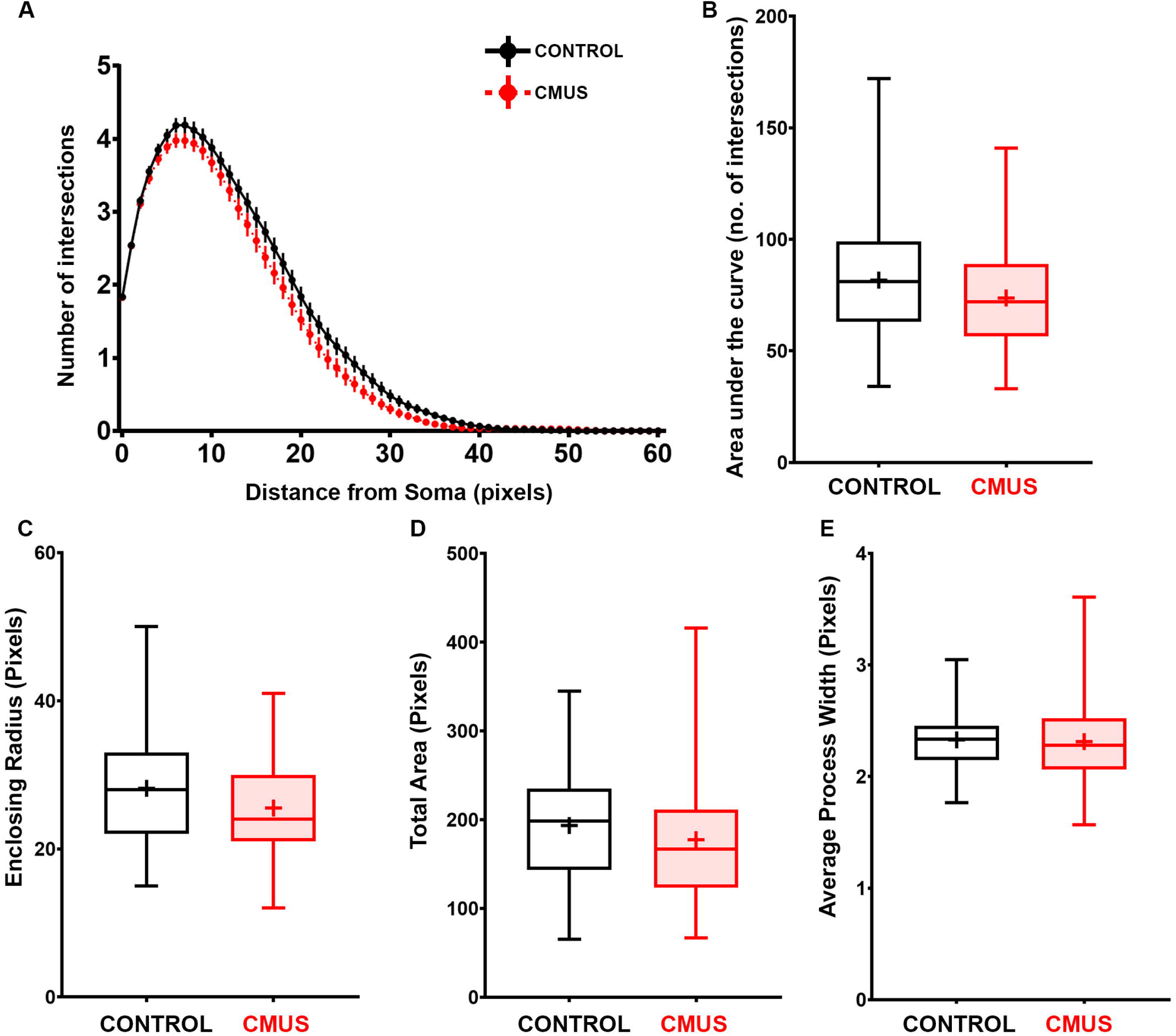
Effects of CMUS on the morphology of astrocytes in the stratum radiatum of CA1. Mice were subjected to 21 days of Chronic Mild Unpredictable Stress (CMUS), while the controls were handled similarly without exposure to stress. GFAP immunostained astrocytes were imaged on a confocal microscope and the images of astrocytes from the *stratum radiatum* region of the CA1 were analyzed for morphological analysis as shown in Figure 4. Line plot depicting Sholl analysis reveals no significant difference in the ramification of astrocytes in CMUS-treated mice as compared to the control mice **(A)**. Additionally, Box and whiskers plots show no significant change in the area under the Sholl curve **(B)**, Enclosing Radius **(C)**, Total projected surface area **(D)** and Average process width **(E)** for astrocytes from CMUS-treated mice as compared to the controls. n= (control: 91 cells from 4 mice, CMUS: 73 cells from 3 mice). Data represented as mean ± SEM for Sholl analysis and as box and whisker’s plot for two group comparisons. Sholl analysis data was analyzed by two-way repeated measures ANOVA followed by Sidak’s *post-hoc* test. Two group comparisons were analyzed with nested t-test.

## Discussion

Hippocampus is an integral part of the limbic system and is implicated in the pathophysiology of MDD; as well as in the action of antidepressant drugs (Nestler et al., 2002). Moreover, astrocytic populations in specific subfields of the hippocampus seem to be differentially vulnerable to the pathology associated with MDD (Cobb et al., 2013, 2016). The quest for the mechanistic understanding of these differential vulnerabilities between different astrocytic subpopulations of the hippocampus necessitates a careful profiling in a tractable model system such as mice. In this study, we used CMUS mouse model and assessed its effects on the cell densities and morphology of hippocampal astrocytes from different subfields (Table 1). We found that the astrocytic density was significantly lowered selectively in the hilus and CA1, specifically in the rostral sections. Furthermore, in terms of morphological characteristics related to atrophy, molecular layer was found to be the most affected subfield, followed by the hilus. CA3 astrocytes were much less affected, while the astrocytes in CA1 were largely spared. These results corroborate the observations in MDD patients showing that the astrocytic subpopulations from different hippocampal subfields are differentially vulnerable to MDD pathology (Cobb et al., 2013, 2016).

**Table 1:**
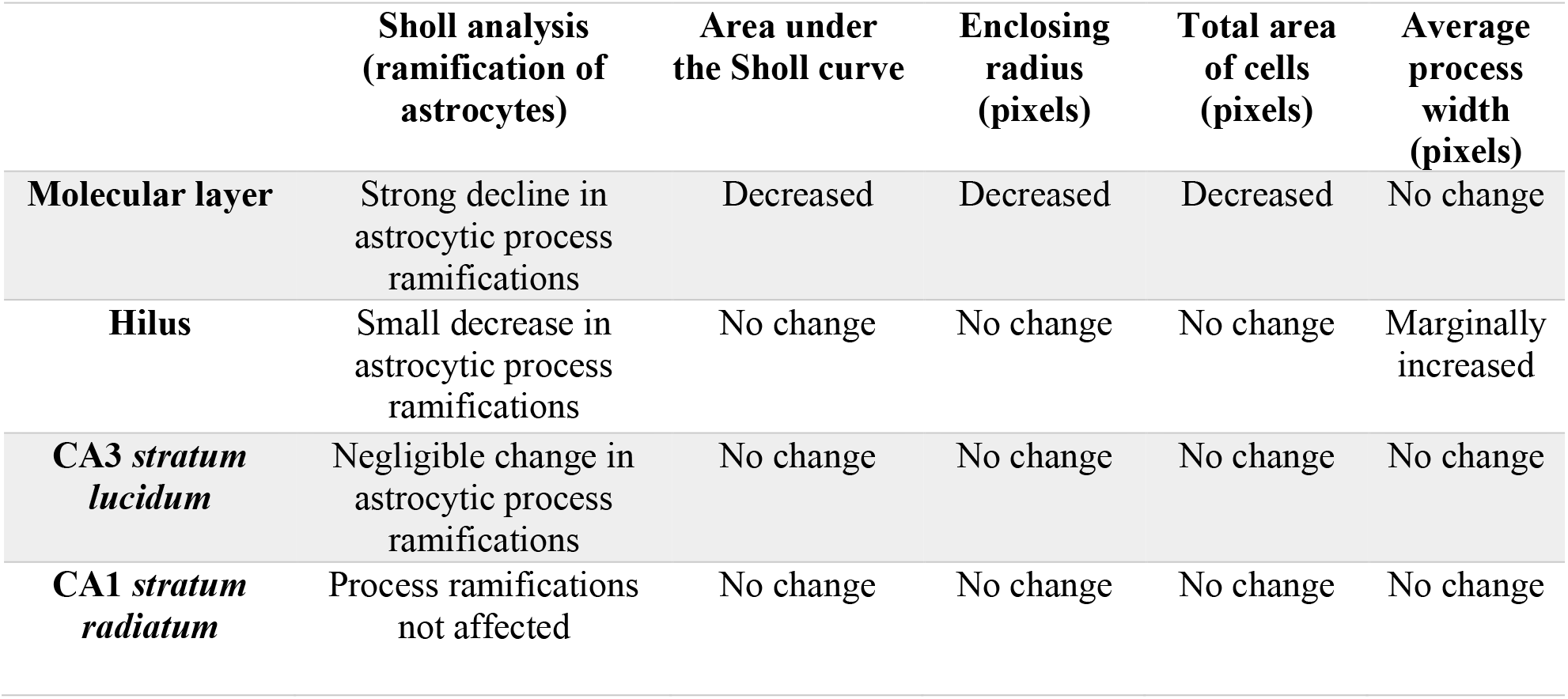
Summary of the effects of CMUS on astrocyte morphology. Table shows the summary of the effects of CMUS on the morphology of hippocampal astrocytes from molecular layer of the DG, Hilus, CA3 *stratum lucidum* and CA1 *stratum radiatum*.

|  | **Sholl analysis**<br>**(ramification of**<br>**astrocytes)** | **Area under**<br>**the Sholl curve** | **Enclosing**<br>**radius**<br>**(pixels)** | **Total area**<br>**of cells**<br>**(pixels)** | **Average**<br>**process**<br>**width**<br>**(pixels)** |
| --- | --- | --- | --- | --- | --- |
| **Molecular layer** | Strong decline in astrocytic process ramifications | Decreased | Decreased | Decreased | No change |
| **Hilus** | Small decrease in astrocytic process ramifications | No change | No change | No change | Marginally increased |
| **CA3 *stratum lucidum*** | Negligible change in astrocytic process ramifications | No change | No change | No change | No change |
| **CA1 *stratum radiatum*** | Process ramifications not affected | No change | No change | No change | No change |

### Analysis of astrocyte morphology in animal models of stress

Studies in animal models have shown divergent results regarding the effects of stress on astrocytes. It was shown that chronic psychosocial stress in treeshrews decreases the number and somal volume of hippocampal astrocytes, which was reversed by chronic fluoxetine treatment (Czéh et al., 2005). On the other hand, studies have also shown no differences in the hippocampal astrocytes between stressed groups and controls (Musholt et al., 2009; Naskar and Chattarji, 2019). Another study also showed an increase in the hippocampal GFAP immunoreactivity after chronic immobilization stress (Jang et al., 2008). However, these differences can arise from differences in experimental stress paradigms and the subfields of the hippocampus that were studied. Consistent with some of these studies, we found no differences in astrocyte morphology in the dorsal CA3 (Naskar and Chattarji, 2019) (Figure 7), as well as no statistically significant differences in the number of astrocytes in CA1, when the sections were taken spanning the rostro-caudal extent (Musholt et al., 2009) (Figure 2). Our study addresses some of these concerns by simultaneously analyzing different hippocampal subfields in the same group of mice. We believe that these results can be useful to further probe the mechanisms underlying differential vulnerabilities to stress between different astrocytic subpopulations.

These studies also raise further questions regarding the effects of stress on astrocytic structure and function. Fine leaflet-like process, also called perisynaptic astrocytic process, ensheath synapses in the central nervous system and play an integral role in the regulation of synaptic transmission (Bernardinelli et al., 2014). However, GFAP only stains about 15% of the GFAP somal volume revealing only the core astrocytic processes. Hence, future studies will need to address the plasticity at the perisynaptic astrocytic processes.

Such studies usually require dye-filling experiments to stain the cytoplasm in its entirety (Medvedev et al., 2014; Naskar and Chattarji, 2019). This makes these experiments time consuming, while severely limiting the ability to analyze large sample sizes across different brain regions. Sparsely labeling the astrocytes *in vivo* with genetically encoded membrane-targeted fluorescent proteins would allow the analysis of large datasets encompassing various brain regions across several experimental paradigms. Automation of some of the steps in the image-processing pipeline would further facilitate these large-scale analyses.

### Differential involvement of dorsal versus ventral hippocampus in pathophysiology of depression

Hippocampus is a large brain structure neatly arranged into a trisynaptic circuitry, which is involved in learning and memory as well as emotional behavior. Interestingly, hippocampus exhibits neuroanatomical (Ruth et al., 1982; Van Groen and Lopes da Silva, 1985; Witter, 1986), molecular (Cembrowski et al., 2016a, 2016b; Lee et al., 2017), electrophysiological (Witter, 1986) as well as functional (Bannerman et al., 2003; Moser and Moser, 1998) dichotomy along its posterio-anterior axis, which corresponds to the dorso-ventral axis in rodents. While the posterior hippocampus, or dorsal hippocampus in rodents is involved in memory encoding (Moser and Moser, 1998), the anterior or ventral hippocampus is involved in processing emotional information (Bannerman et al., 2003). Supporting this view of functional segregation, it was shown that astrocyte cell density in the anterior hippocampus was decreased in depressed female monkeys (Willard et al., 2013). In this study as well, only the astrocyte density in CA1 and DG was selectively diminished (Willard et al., 2013). Intriguingly, we found that the astrocyte density is lower in stressed mice in the rostral (Bregma −1.46 to −1.94) coronal sections, which correspond to the dorsal hippocampus, while we saw that in the sections from more caudal (ventral) sections, astrocyte density remained unchanged. This is indeed interesting, given that ventral hippocampus receives denser innervations from the hypothalamus, the master regulator of stress responses. That said, the caudal coronal sections (Bregma −2.54 to −2.92) from rodent brain are not entirely ventral hippocampal but have sizable portions from the intermediate hippocampus. Transverse sections of dissected hippocampi allow a *bona-fide* distinction between the dorsal and ventral poles of the hippocampus. Further studies are needed to ascertain whether astrocyte density in the ventral hippocampus is indeed unaltered in CMUS-treated mice.

### Astrocytic atrophy as a potential pathogenic event

Psychiatric and neurodegenerative disorders can be viewed as synaptopathies (Brose et al., 2010). Astrocytic atrophy and dysfunction has been reported in several neurological and psychiatric disorders; including MDD (Cotter et al., 2001; Miguel-Hidalgo et al., 2000; Ongür et al., 1998; Rajkowska and Stockmeier, 2013; Rajkowska et al., 1999; Verkhratsky et al., 2014). Atrophy in astrocytes leads to reduction in synaptic coverage thereby compromising the wellbeing of synapses (Verkhratsky et al., 2016). This phenomenon may manifest itself through reduced glutamate uptake, diminished trophic support or disruption of metabolic coupling (Elsayed and Magistretti, 2015; Marathe et al., 2018; Martin et al., 2013). Despite such a consequential nature of astroglial atrophy in several brain disorders, the molecular mechanisms underlying the atrophy remain elusive. It was shown that selective ablation of prefrontal astrocytes in rats was sufficient to induce depressive-like symptoms in experimental rats (Banasr and Duman, 2008). Hence, it is conceivable that preventing the glial degeneration associated with MDD may prove to be a viable therapeutic strategy.

### Conclusions

In conclusion, while majority of MDD research so far has focused on neurons, the astrocytic degeneration and dysfunction has remained relatively unexplored. The data from this manuscript highlights the fact that not all astrocytes within the hippocampus are uniformly affected by stress. This corroborates previous studies involving postmortem analyses in the victims of MDD showing differential vulnerabilities within hippocampal subfields (Cobb et al., 2013, 2016). Given the integral role of hippocampus in MDD pathophysiology and in antidepressant action, astrocytic degeneration in selective subfields of hippocampus may have important functional ramifications.

## Acknowledgements

This work was supported by INSPIRE faculty grant from Department of Science and Technology (DST), India to SM and Early Career Research Award from Scientific and Engineering Research Board (SERB) to SM. GV and AN were supported by CSIR-NET Junior and Senior Research fellowship. Authors are grateful to Prof. Narendrakumar Ramanan for S100β antibody and critical comments on this manuscript. Authors would also like to thank Mr. Manjunath and the staff at central animal facility, IISc and bioimaging facility, IISc.

## Conflict of interest statement

The authors declare that the research was conducted in the absence of any commercial or financial relationships that could be construed as a potential conflict of interest.

## Author contributions

SM conceived and designed the study. PD, GV and SM performed experiments. GV analyzed the data. AN performed FST behavior analysis. SM wrote the manuscript. All authors discussed, proofread and approved the final version.

## Data Accessibility Statement

Requests for raw data can be addressed to the corresponding author and the data will be made available upon reasonable request.

## Abbreviations

BSA: Bovine Serum Albumin
CA3 s.l.: *stratum lucidum* region of CA3
CA1 s.r.: *stratum radiatum* region of CA1
CMUS: Chronic Mild Unpredictable Stress
DG: Dentate Gyrus
FST: Forced Swim Test
MDD: Major Depressive Disorder
PB: 0.1M Phosphate Buffer
PBS: Phosphate Buffered Saline
PFA: Paraformaldehyde
ROI: Region of Interest.

