## Supplemental Information for "Subfield-specific Effects of Chronic Mild Unpredictable Stress on Hippocampal Astrocytes"

**Supplementary Table 1**

| | First Stressor | Second Stressor |
|---|---|---|
| **Day 0** | Forced Swim Stress | |
| **Day 1** | Overcrowding | Cage Tilt |
| **Day 2** | Tail Suspension Stress | Cold Exposure |
| **Day 3** | Overcrowding | Cage Tilt |
| **Day 4** | Wet Bedding | Restraint |
| **Day 5** | Forced Swim Stress | Food Water Deprivation |
| **Day 6** | Cold Exposure | |
| **Day 7** | Tail Suspension Stress | |
| **Day 8** | Wet Bedding | Cage Tilt |
| **Day 9** | Shaker Stress | Tone |
| **Day 10** | Forced Swim Stress | White Noise |
| **Day 11** | Overcrowding | Tone |
| **Day 12** | Tail Suspension Stress | Cold Exposure |
| **Day 13** | Restraint | |
| **Day 14** | Wet Bedding | |
| **Day 15** | Forced Swim Stress | Cage Tilt |
| **Day 16** | Overcrowding | Tone |
| **Day 17** | Tail Suspension Stress | Wet Bedding |
| **Day 18** | Shaker Stress | Food Water Deprivation |
| **Day 19** | Forced Swim Stress | Cage Tilt |
| **Day 20** | Overcrowding | |
| **Day 21** | Food Water Deprivation | |
| **Day 22** | Sacrifice | |

**Supplementary Figure 1**

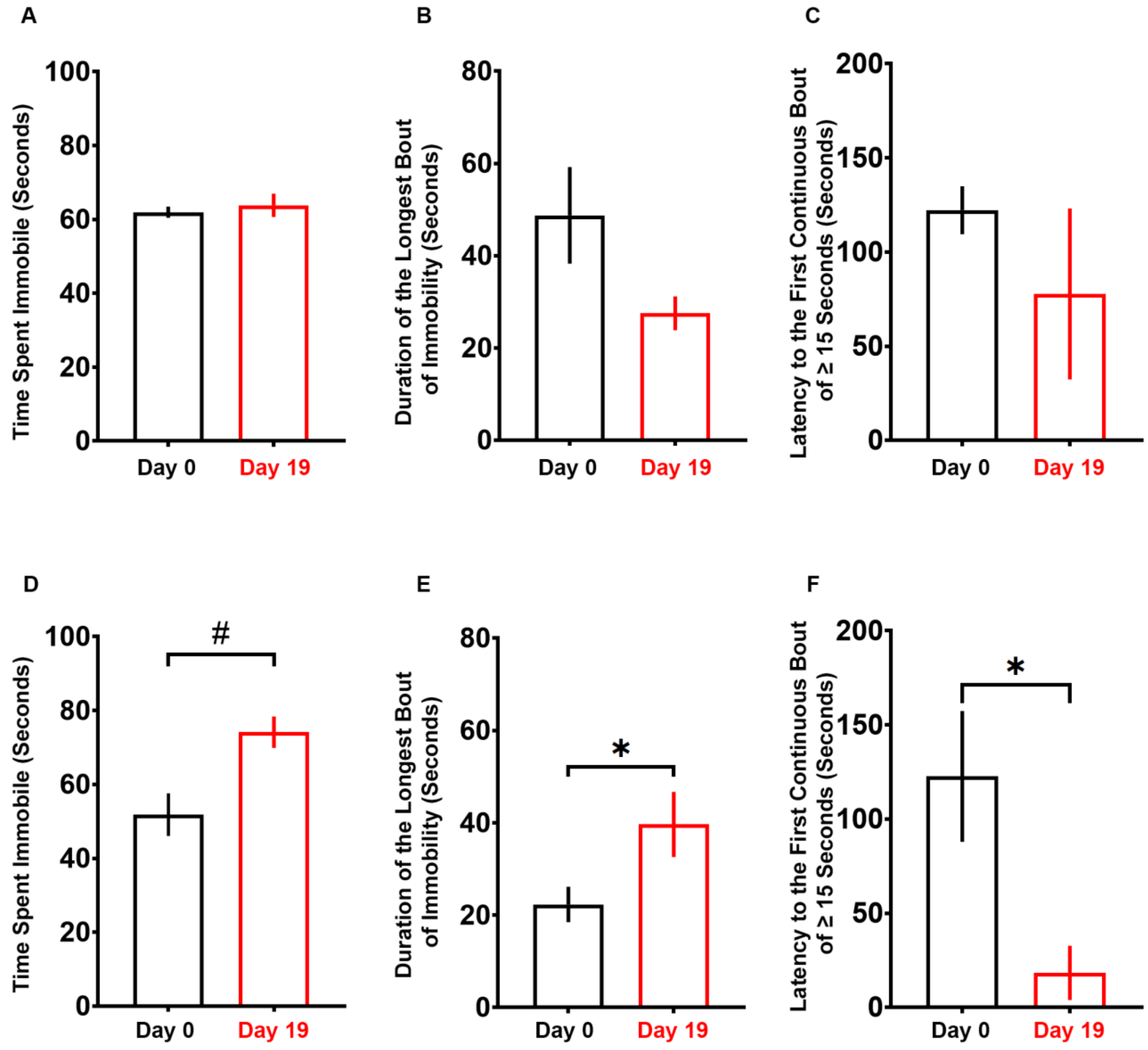

**Supplementary Figure 2**

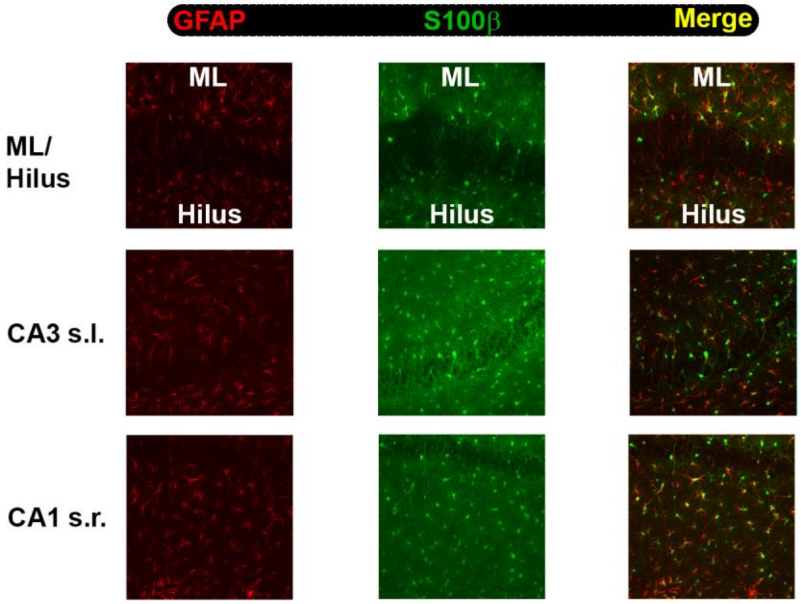

**Supplementary Figure and Table Captions:**

*Supplementary Figure 1*

Mice were subjected to 21 days of Chronic Mild Unpredictable Stress (CMUS), while the controls were handled similarly without exposure to stress. The depressive-like behavior was monitored using Forced Swim Test (FST). The control mice did not show any change in the time spent immobile between day 0 and day 19 **(A)**. The control mice also did not show any change in the duration of the longest bout of immobility and the latency to a continuous bout of immobility of 15 seconds or longer **(B, C)**. On the other hand, the CMUS-treated mice showed a trend towards an increase in time spent immobile from day 0 to day 19 **(D)**. The CMUS-treated mice also showed an increase in the duration of the longest bout of immobility **(E)** and a decrease in the latency to a continuous bout of immobility of 15 seconds or longer **(F)**. n=3-4 mice per group. Data represented as mean ± SEM. # represents $0.05<p<0.1$. * represents $p < 0.05$. All comparisons made using paired student's t-test.

*Supplementary Figure 2*

The sections were double immunolabeled with GFAP and S100β and confocal images were obtained. Representative maximum intensity projection images of molecular layer (ML) of the dentate gyrus, Hilus, *stratum lucidum* layer of CA3 (CA3 s.l.) and *stratum radiatum* region of CA1 (CA1 s.r.) showing immunostaining for GFAP (red), S100β (green) and Merged channels showing a near complete overlap between cells expressing GFAP and S100β.

*Supplementary Table 1*

Table shows the schedule of different stressors employed during the 21-day CMUS paradigm. On day 22, 24 hours after the last stressor, mice were sacrificed.
